# Growth-dependent tRNA Reprogramming and Codon Bias Link Translation to Metabolic State in *Enterococcus faecalis*

**DOI:** 10.64898/2026.05.07.723122

**Authors:** Michelle M. Mitchener, Caleb M. Anderson, Mark Veleba, Kayla Teo, Mélanie Roch, Ruixi Chen, Yifeng Yuan, Isaiah Han, Agnieszka Dziergowska, Kevin Pethe, Thomas J. Begley, Kimberly A. Kline, Peter C. Dedon

**Affiliations:** Antimicrobial Resistance Interdisciplinary Research Group, Singapore-MIT Alliance for Research and Technology Centre, Singapore 138602, Singapore; Department of Microbiology and Molecular Medicine, Faculty of Medicine, University of Geneva, 1211 Geneva 4, Switzerland; Singapore Centre for Environmental Life Sciences Engineering, Nanyang Technological University, Singapore 637551, Singapore; Department of Biological Engineering, Massachusetts Institute of Technology, Cambridge, MA 02139, USA; Institute of Organic Chemistry, Lodz University of Technology, 90-924 Lodz, Poland; Lee Kong Chian School of Medicine, Nanyang Technological University, Singapore 636921, Singapore; National Centre for Infectious Diseases (NCID), Singapore 308442, Singapore; Department of Biological Sciences, University at Albany, Albany, NY 12222, USA; RNA Institute, University at Albany, Albany, NY 12222, USA; School of Biological Sciences, Nanyang Technological University, Singapore 637551, Singapore

## Abstract

*Enterococcus faecalis* is a Gram-positive commensal bacterium of the human gut microbiome and an opportunistic pathogen responsible for many hospital-acquired infections. Despite the clinical importance of *E. faecalis*, how gene and protein expression are coordinated with growth remains poorly defined. Here, we profiled transcript, protein, and tRNA pool dynamics across distinct phases of *E. faecalis* growth. Differences in protein abundance and corresponding mRNA levels suggested growth phase-dependent posttranscriptional regulation. Growth-associated genes exhibited biased synonymous codon usage, with ribosomal and glycolytic proteins enriched in low-abundance codons read by queuosine-modifiable tRNAs. Analysis of tRNA modification and tRNA isoacceptor abundance revealed growth phase-dependent changes, particularly in anticodon stem loop modifications that influence synonymous codon translation. Changes in queuosine levels preceded shifts in ribosomal proteins, suggesting a contribution to codon-biased translation. Collectively, these findings reveal growth phase-associated remodeling of the *E. faecalis* tRNA pool and support a model in which queuosine-dependent translational reprogramming shapes protein expression during bacterial growth.

**IMPORTANCE:** *Enterococcus faecalis* is a common cause of hospital-acquired infections. Despite its clinical importance, a comprehensive understanding of the organism’s physiology and adaptation to environmental changes remains incomplete. Here, we characterized protein, transcript, and tRNA dynamics across bacterial growth phases, uncovering a role for post-transcriptional regulation marked by tRNA reprogramming and biased synonymous codon usage. These findings enhance our understanding of *E. faecalis* growth and support a model of translational reprogramming therein.

## INTRODUCTION

*Enterococcus faecalis* is a common inhabitant of the human gastrointestinal tract than can be opportunistically pathogenic when mucosal immunity is compromised, the gastrointestinal barrier is disrupted, or the native microbiota is depleted following antibiotic treatment (1, 2). Consequently, *E. faecalis* is a frequent cause of hospital-acquired infections, including central-line associated bloodstream infections, catheter-associated urinary tract infections, surgical site infections, and infective endocarditis (3, 4). Treatment is often complicated by antibiotic resistance (5, 6). Moreover, many *E. faecalis* are biofilm-associated (7), giving rise to phenotypic antibiotic tolerance, underscoring the need for new therapeutic strategies (8).

Despite this clinical imperative, the regulatory mechanisms that coordinate gene expression with growth and environmental adaptation in *E. faecalis* are incompletely understood. Bacteria must rapidly adjust protein synthesis in response to changes in temperature, pH, oxygen availability, nutrient levels, antibiotic exposure, and competition with other microorganisms. One mechanism that facilitates such rapid adaptation is remodeling of the tRNA pool, comprising both tRNA isoacceptors and their modifications. Changes in tRNA abundance or modification can promote selective translation of mRNA transcripts enriched in specific synonymous codons, thereby biasing protein synthesis toward transcripts that support distinct physiologic states (9, 10). Such codon-biased translation has been linked to cellular responses to growth conditions (11–20), oxidative and chemical stress (18, 19, 21–25), and antibiotic exposure (26–29). These dynamics arise through coordinated changes in the expression or activity of tRNA-modifying enzymes, as well as in the synthesis and degradation of tRNAs. Consistent with their regulatory importance, disruption of tRNA-modifying enzymes can impair bacterial fitness (30–33), sporulation (34, 35), biofilm formation (34), intracellular replication (19, 32), and host colonization (36–41). Whether tRNA pool remodeling contributes to growth-dependent translational regulation in *E. faecalis* remains unknown.

Here, we examined *E. faecalis* OG1RF growth by quantifying transcripts, proteins, tRNA modifications, and tRNA isoacceptors across distinct phases of bacterial growth (**Fig. 1**). Discordant changes between mRNA and protein abundance over time suggested growth-dependent posttranscriptional regulation. Analysis of protein-coding genes revealed strong codon biases in growth-associated proteins, particularly ribosomal proteins, which preferentially use C-ending codons read by queuosine (Q)-modifiable tRNAs. Moreover, tRNA modification and isoacceptor abundance changed dynamically during growth, with Q modification shifts preceding similar trends in ribosomal protein abundance. Collectively, these data provide an integrated understanding of *E. faecalis* growth and support a role for tRNA reprogramming and codon-biased translation in this process.

**Fig. 1.**
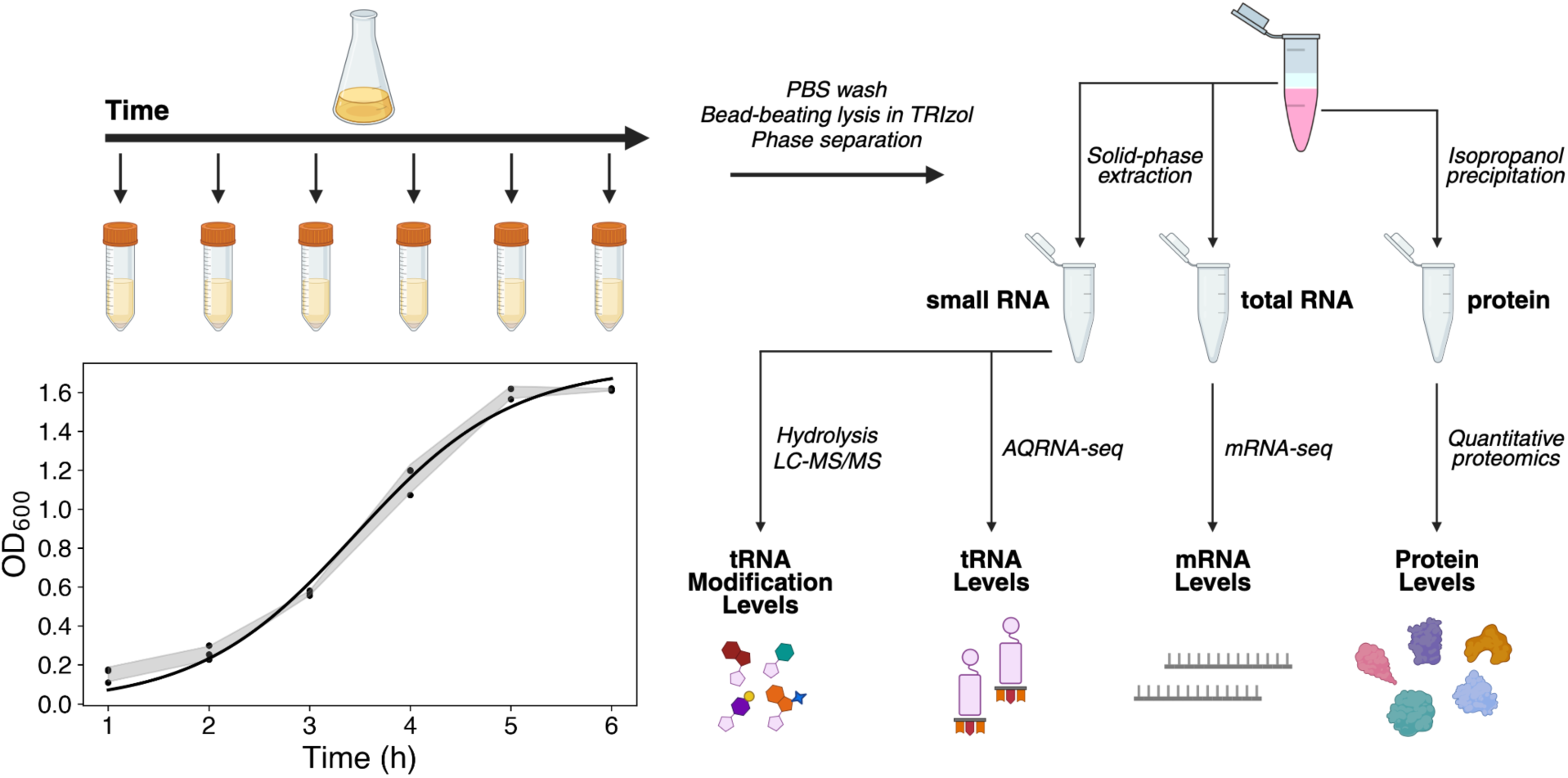
Experimental approach to characterize *E. faecalis* growth. Biological triplicate samples were taken over six hours, with spectrophotometric monitoring of cell density (bottom left, shaded region represents ± SD, sigmoidal curve fit). Analytes were isolated and quantified as indicated.

## RESULTS

### Protein abundance during E. faecalis growth is not solely explained by transcription

We began by examining protein abundance across a six-hour *E. faecalis* growth time course. Using TMT-based quantitative proteomics, we detected 1627 proteins, representing 65% proteome coverage (**Table S1**). Proteins with similar temporal patterns were grouped using fuzzy clustering (42), yielding eight distinct clusters (**Fig. 2A** and **Table S2**) enriched for specific KEGG and GO pathways (**Fig. 2B** and **Table S3**). Proteins involved in ATP biosynthesis, proton transmembrane transport, lysine biosynthesis, and pyrimidine metabolism (Cluster 3) decreased through log phase and then increased as cells approached stationary phase, suggesting a renewed demand for these pathways during slow growth. Carbohydrate transport proteins and associated phosphotransferase machinery (Cluster 7) declined rapidly upon entry into log phase and gradually increased thereafter, consistent with reduced uptake requirement during nutrient abundance followed by increased expression as resources become limiting. By contrast, glycolytic enzymes, ABC transporters, and ribosomal proteins (Clusters 4, 6, and 8, respectively) increased from lag to log phase, consistent with elevated metabolic and translational demands during rapid cell growth. Whereas glycolytic proteins remained elevated in stationary phase, levels of ABC transporters and particularly ribosomal proteins declined, consistent with reduced nutrient uptake and translational activity.

**Fig. 2.**
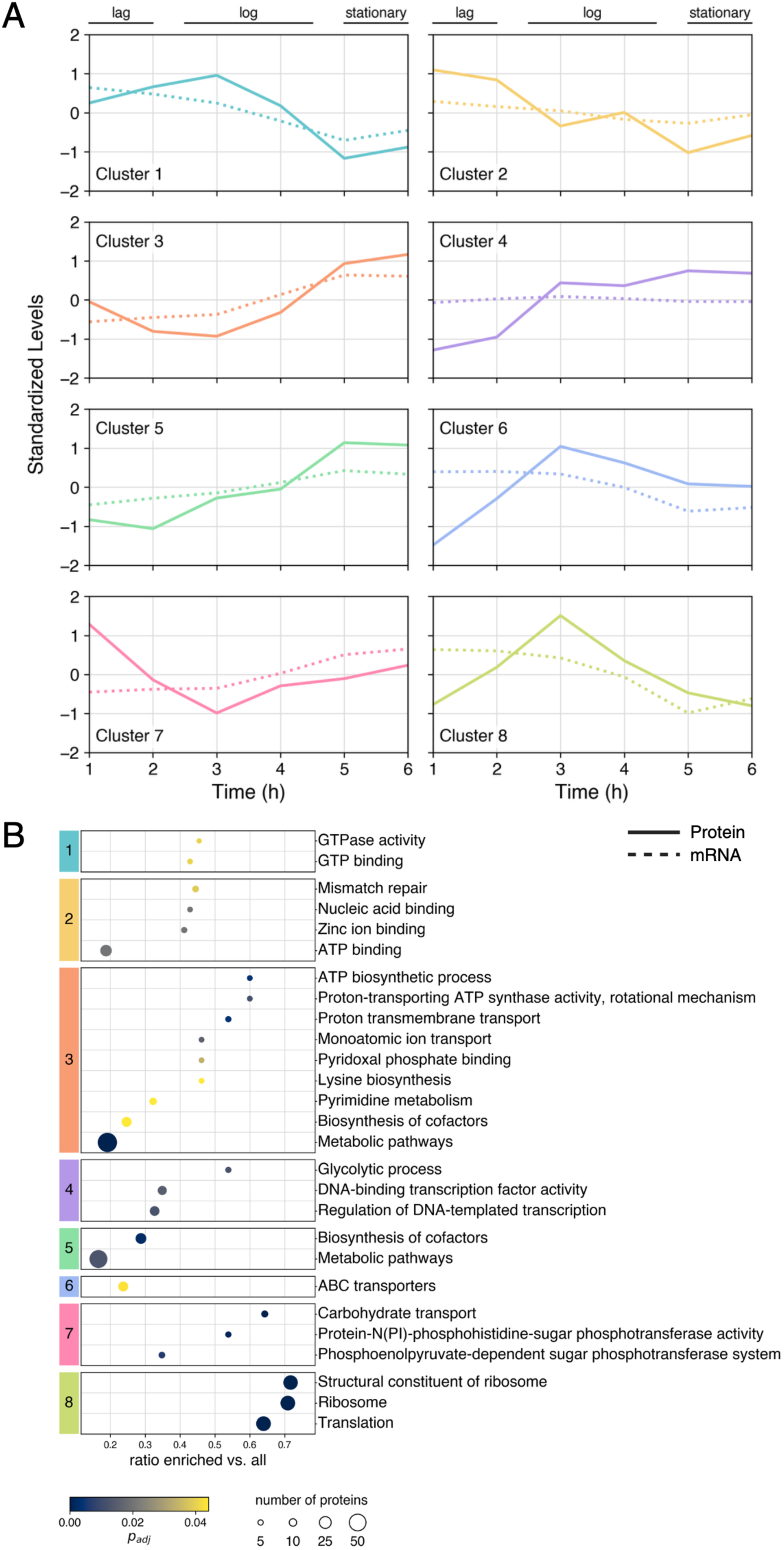
Discordant trends observed between protein and corresponding transcript levels. **(A)** Fuzzy clustering analysis of quantitative proteomics data over time, with protein centroids shown as solid lines and mRNA centroids shown as dashed lines (n = 3 biological replicates). **(B)** Overrepresentation analysis of each cluster, with enriched KEGG and GO pathways indicated (p_adj_ < 0.05).

To determine the extent to which protein levels were transcriptionally driven, we quantified mRNA abundance over the same time course using RNA-seq (**Fig. 2A** and **Table S4**). In some clusters (e.g. Clusters 2 and 5), mRNA and protein profiles were similar suggesting dominant transcriptional control. In others (especially Clusters 4, 6, 7, and 8), transcript and protein trajectories diverged, particularly at early time points. Such discordance is commonly observed and suggests the involvement of posttranscriptional regulatory mechanisms (43, 44). This raised the possibility that tRNA reprogramming and associated codon-biased translational regulation contribute to protein expression during *E. faecalis* growth.

### Genes encoding growth-associated proteins exhibit distinct synonymous codon biases

If tRNA reprogramming and codon-biased translation contributes to posttranscriptional regulation in *E. faecalis*, genes encoding growth-associated proteins would be expected to exhibit biased synonymous codon usage. To test this possibility, we calculated codon usage and isoacceptor codon frequencies of all *E. faecalis* OG1RF protein-coding genes and performed Principal Component Analysis (PCA) on the corresponding z-score data (Fig. 3A and **B**; **Table S5**) (9). Although many genes cluster closely in two-dimensional PCA space, a subset differ substantially in their codon usage compared to the rest of the genome. Overlaying KEGG and GO pathway annotations onto the PCA plot revealed that many growth-associated proteins occupy distinct regions of codon usage space (**Fig. 3A**). The corresponding loadings plot identified the synonymous codons contributing most strongly to this separation (**Fig. 3B**). Among these were codons for amino acids Asn, Asp, His, and Tyr, which are each encoded by two synonymous codons and decoded by single, modifiable GUN anticodons on their corresponding tRNAs (**Table S6**). Glycolytic and ribosomal proteins preferentially use C-ending Asn, Asp, His, and Tyr codons compared to their T-ending (U-ending in mRNA) counterparts (**Fig. 3C** through **F**). Ribosomal proteins in particular are strongly enriched in C-ending Asn and Tyr codons, and to a lesser extent, C-ending Asp and His codons, compared to the genome-wide median. Moreover, these C-ending codons are underrepresented in the genome, occurring nearly three times less frequently than their T/U-ending counterparts (**Fig. 3C** through **F**; **Table S5**). T/U-ending codons predominate in proteins important for bacteria survival in reduced-glucose environments, including phosphoenolpyruvate-dependent sugar phosphotransferase system and carbohydrate transport proteins. This preferential use of synonymous codons in growth-associated proteins suggested that their translation might be subject to tRNA reprogramming-based regulation.

**Fig. 3.**
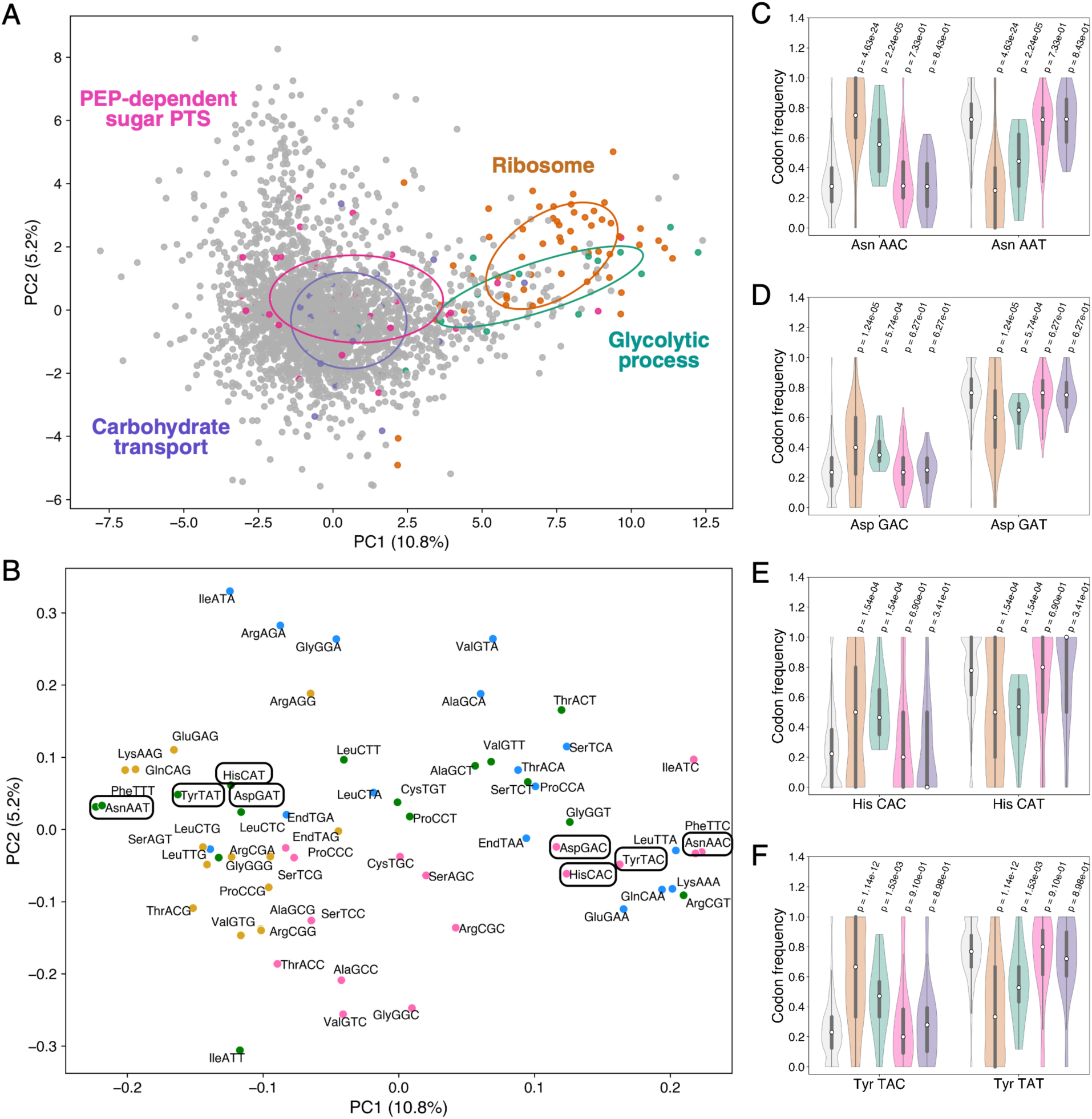
Growth-associated pathway proteins are codon-biased. **(A)** Principal component analysis (PCA) scores plot based on *E. faecalis* OG1RF protein-encoding gene codon usage z-scores. Ribosome, glycolytic process, phosphoenol (PEP)-dependent sugar phosphotransferase system (PTS), and carbohydrate transport pathways are indicated in orange, teal, pink, and purple, respectively. Covariance ellipses are also depicted in pathway-specific colors. **(B)** PCA loadings plot with codons colored according to terminal nucleotide (A: blue; T: green; G: yellow; C: pink). Codons read by Q-bearing tRNA isoacceptors are circled. Violin plot of the frequency of codons highlighted in **(B)** in proteins from the KEGG/GO pathways shown in **(A)**: ribosome (**(C)**, orange)), glycolytic process (**(D)**, teal), PEP-dependent sugar PTS (**(E)**, pink), and carbohydrate transport (**(F)**, purple) versus in the entire genome (gray). Violin width represents density, thick gray bars the interquartile range (IQR), thin lines 1.5xIQR, and white dots the median. P-values were determined by Mann-Whitney U testing with Benjamini-Hochberg FDR correction.

### tRNA modifications exhibit distinct growth-phase behaviors

Since tRNA reprogramming involves changes in both tRNA modification and tRNA isoacceptor abundance, we next characterized each across the *E. faecalis* growth curve. Purified small RNA (> 85% tRNA) was digested to ribonucleosides and modification levels were quantified by liquid chromatography-coupled tandem mass spectrometry (LC-MS/MS) (**Table S7**). Several modifications, including queuosine (Q), 5-methoxyuridine (mo^5^U), 4-thiouridine (s^4^U), and 2-methyladenosine (m^2^A), displayed pronounced growth-dependent dynamics (Fig. 4A and **B**).

**Fig. 4.**
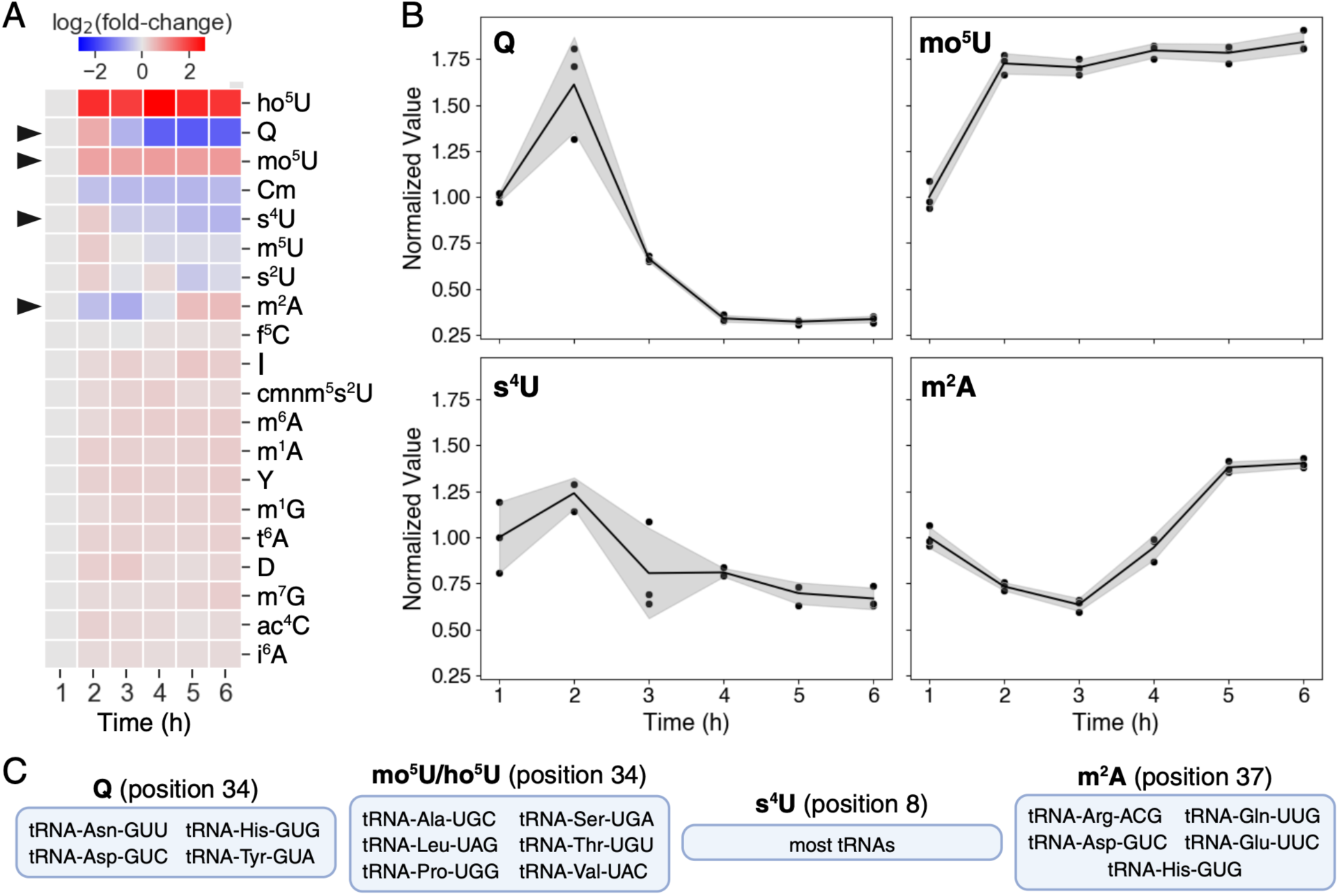
tRNA modification changes during growth. **(A)** Heatmap of small RNA modification levels over the growth time-course, expressed as log_2_(fold-change) with respect to modification levels at t = 1 h. Values represent the average of three biological replicates. **(B)** Scatterplot of select dynamic modifications (indicated by arrows in **(A)**) over time, expressed relative to modification levels at t = 1 h. Shaded regions represent the mean ± SD. **(C)** tRNA isoacceptors and the nucleotide positions therein on which the modifications in **(B)** have been reported (45).

Q levels increased sharply in late lag phase and then decreased during log and stationary phases (Fig. 4A and **B**). Q is found at position 34, the first nucleotide of the anticodon, in Asn, Asp, His, and Tyr tRNAs, where it replaces G in their respective GUN anticodons (**Fig. 4C** and **Fig. S1A**) (45). Homology analysis suggests that Q is installed into *E. faecalis* tRNA through the sequential action of three enzymes: Tgt, QueA, and QueG (**Fig. S1B**) (46). Consistent with Q modification dynamics, Tgt protein levels decreased sharply during log phase (3 h) (**Fig. S1C**). Correlations with QueA and QueG were more nuanced; however, QueG levels increased during lag phase and decreased after log phase (**Fig. S1D** and **E**). These trends are consistent with the detection of the preQ_1_ modification only at the 1 h time point, suggesting conversion to Q by late lag phase followed by a subsequent decline (**Table S7**). Previous studies have linked Q levels to reactive oxygen species (ROS), suggesting that increased Q may occur under oxidative conditions (25, 47, 48). Consistent with earlier reports (49), we observed elevated H_2_O_2_ during *E. faecalis* lag phase that decreased through stationary phase in parallel with Q changes (**Fig. S2**). This observation raises the possibility that the transient Q spike is triggered by ROS accumulation early in growth.

Levels of mo^5^U and its precursor, 5-hydroxyuridine (ho^5^U), increased during lag phase and remained elevated over the time course (Fig. 4A and **B**). These modifications occur at position 34 in select Ala, Leu, Pro, Ser, Thr, and Val tRNAs (**Fig. 4C** and **Fig. S3A**) (45). ho^5^U is synthesized from uridine by TrhP1/TrhP2 and/or TrhO, which use prephenate and molecular oxygen as the hydroxyl source, respectively (**Fig. S3B**) (50). ho^5^U is then methylated to mo^5^U by the S-adenosylmethionine (SAM) methyltransferase TrmR (51). TrhP1 and TrhP2 homologs increased during lag phase, after which TrhP1 decreased slightly while TrhP2 continued to rise through log phase before stabilizing (**Fig. S3C** and **D**). These trends are consistent with the observed increase in ho^5^U in lag phase followed by a plateau in modification levels (**Fig. 4B**). TrmR abundance remained largely unchanged over the time course (**Fig. S3E**), suggesting that substrate availability may limit mo^5^U formation (Fig. 4A and **B**).

s^4^U levels decreased from lag through stationary phase (Fig. 4A and **B**). This modification is present at position 8 between the 5’-acceptor stem and the D-loop in nearly all tRNAs (**Fig. 4C** and **Fig. S4A**) (45). Two enzymes, IscS and ThiI, carry out the coordinated transfer of sulfur from a donor Cys to form s^4^U in tRNA (**Fig. S4B**) (52–54). IscS1 levels decreased from lag through log phase (**Fig. S4C**), while ThiI levels increased up to mid-log phase and then decreased thereafter (**Fig. S4D**). These changes are consistent with the overall decrease in s^4^U from lag through stationary phase (Fig. 4A and **B**).

Levels of m^2^A decreased from lag through log phase and then increased thereafter (Fig. 4A and **B**). This anticodon-adjacent modification occurs at position 37 in select Arg, Asp, Gln, Glu, and His tRNAs (**Fig. 4C** and **Fig. S5A**) (45). The dual-function enzyme RlmN installs m^2^A in both tRNA and rRNA in *E. faecalis* (**Fig. S5B**) (22, 55). Levels of RlmN increased from lag to log phase and then decreased from log to stationary phase, roughly the inverse of the observed m^2^A modification pattern (**Fig. S5C**). Two non-mutually exclusive explanations may account for this discordance between tRNA modification and RlmN protein levels. First, the increase in RlmN abundance parallels the increase in m^2^A levels in 16S/23S rRNA over the time course (**Fig. S6**), consistent with its dual substrate specificity. Second, we previously showed that ROS-inducing antibiotics reduce m^2^A levels in *E. faecalis*, likely via ROS-mediated inactivation of the Fe-S cluster required for RlmN activity (22). Posttranslational inactivation of RlmN by ROS during lag phase (**Fig. S2**) may therefore stimulate compensatory enzyme synthesis until oxidative stress subsides.

### tRNA isoacceptor abundance does not fully explain modification dynamics

To determine whether changes in modification levels reflected altered tRNA abundance, we quantified tRNA isoacceptor abundance using AQRNA-seq (56), a sequencing method optimized for highly modified small RNAs (**Fig. 5A** and **Table S8**). Although some modification dynamics could be rationalized by changes in levels of the tRNAs bearing them, Q was a prominent counterexample (**Fig. 5B**). Q levels increased during late lag phase (2 h) without detectable changes in the abundance of the corresponding tRNA isoacceptors. Levels of Q then declined from log through early stationary phase, while Q-modifiable tRNA-Asn and tRNA-Tyr isoacceptor levels increased. These dynamics could reflect a combination of changes in levels or activity of Q writers and cryptic erasers, degradation of Q-modified tRNAs, and synthesis of unmodified tRNAs. Regardless of the mechanism, these data indicate that the fraction of Q-modified tRNAs increases prior to log phase and decreases thereafter, with unmodified tRNA-Asn and tRNA-Tyr predominating in stationary phase.

**Fig. 5.**
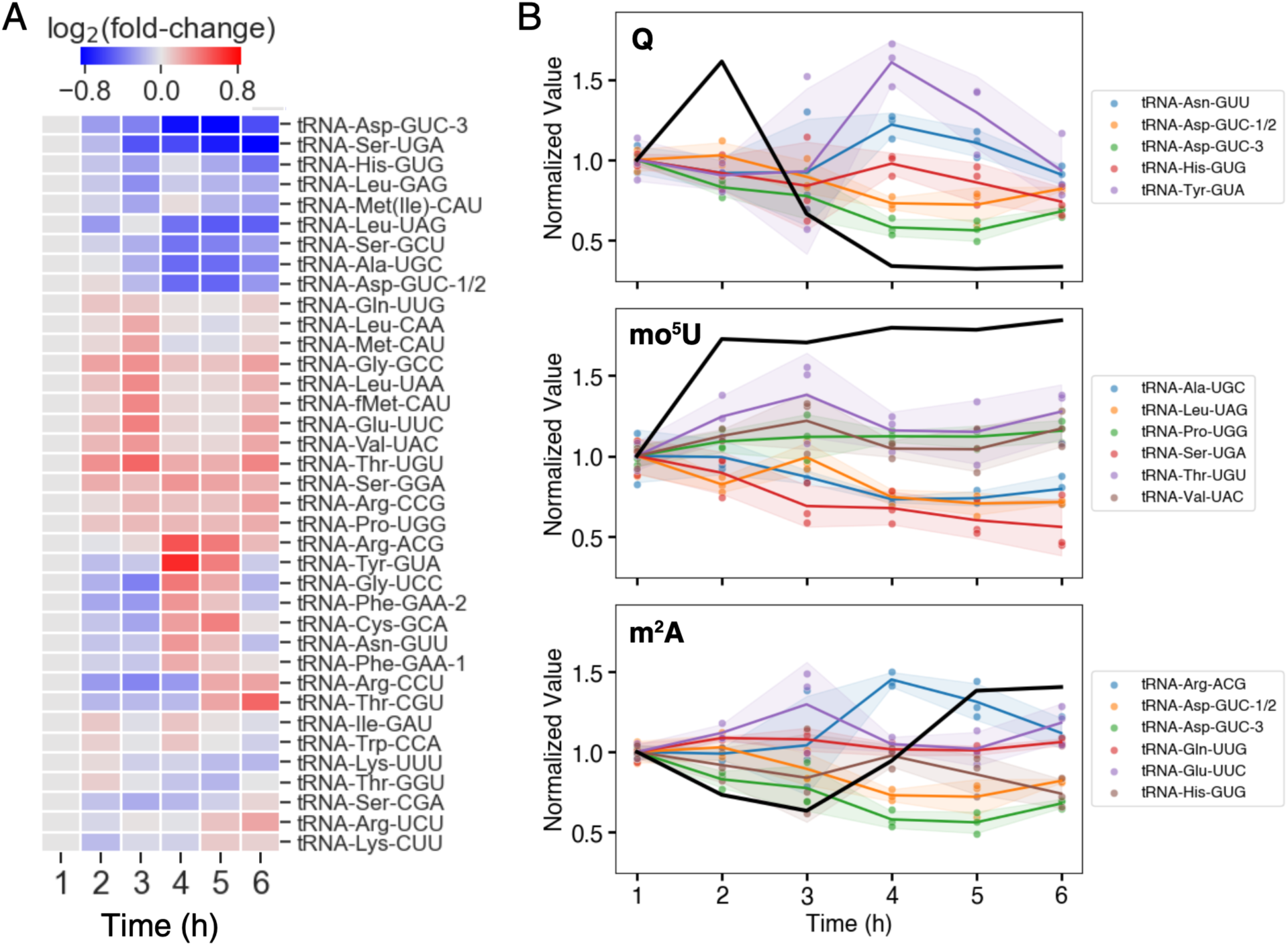
tRNA isoacceptor level dynamics during growth. **(A)** Heatmap of tRNA isoacceptor levels over the growth time-course, expressed as log_2_(fold-change) with respect to tRNA isoacceptor levels at t = 1 h. Values represent the average of three biological replicates. **(B)** Select tRNA modification levels (black lines, from Fig. 4B) overlaid on their reported tRNA isoacceptor levels over the same time-course, expressed relative to the t = 1 h timepoint. s^4^U was not included as it is present on almost all tRNAs. Shaded regions represent the mean ± SD.

## DISCUSSION

The public health burden of *Enterococcus faecalis* and the need for new therapeutic approaches necessitates a comprehensive understanding of the organism’s physiology. A growing body of literature reveals that tRNA reprogramming and codon-biased translation is a common regulatory mechanism in bacteria, but whether it occurs in *E. faecalis* is unknown. Therefore, we undertook a comprehensive characterization of *E. faecalis* growth that identified (1) changes in protein levels that were independent of their corresponding transcripts; (2) biased synonymous codon usage among growth-associated proteins; and (3) growth phase-dependent tRNA pool remodeling. These observations are consistent with a model in which the bacteria respond to environmental cues and modify their tRNAs to promote progression through various growth phases (**Fig. 6**).

**Fig. 6.**
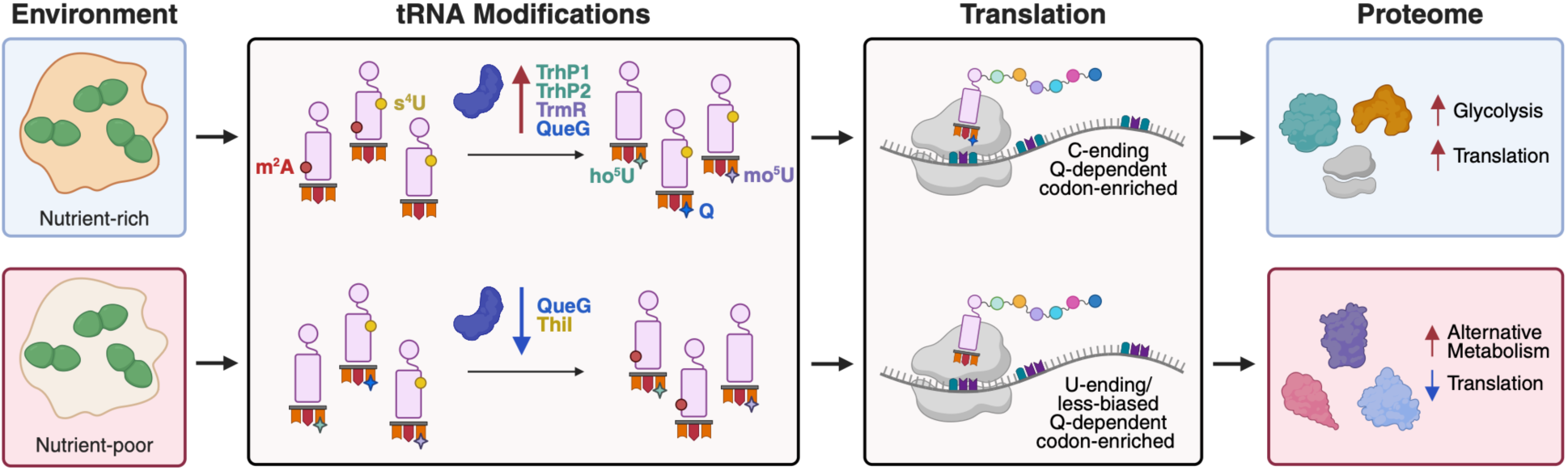
Proposed model of how tRNA reprogramming contributes to *E. faecalis* growth. In nutrient-rich lag phase, increased expression of tRNA-modifying enzymes TrhP1, TrhP2, TrmR, and QueG result in elevated ho^5^U, mo^5^U, and Q levels. These modifications support overall translational efficiency, specifically of C-ending Q-dependent codon-enriched transcripts encoding glycolytic and ribosomal proteins, which characterizes log-phase bacterial growth. In nutrient-poor late log phase, decreased expression of tRNA-modifying enzymes QueG and ThiI result in decreased levels of Q and s^4^U. Lack of these modifications results in reduced overall translational efficiency, as well as the preferential translation of alternative metabolism-associated U-ending Q-dependent codon-enriched transcripts, which characterize stationary-phase bacterial growth.

During nutrient-rich lag phase, Q-modified tRNA levels rise in *E. faecalis*. The translational consequences of Q modification appear complex, with studies reporting preferential translation of either U-ending or C-ending codons, depending on the organism, amino acid, and sequence context (18, 27, 34, 57–61). Here, Q dynamics precede similar trends in glycolytic and ribosomal proteins, which increase through log phase and then decrease thereafter. These proteins are enriched in C-ending Q-dependent codons, suggesting that Q may favor translation of these codons early in *E. faecalis* growth. These protein trends are consistent with a prioritization of translational capacity and glycolytic flux during early growth when preferred carbon sources like glucose are abundant. *E. faecalis* is primarily fermentative and uses this seemingly wasteful strategy (i.e. lower ATP yield per glucose molecule than respiration) to allow rapidly growing bacteria to optimize proteome efficiency, or energy produced per unit protein mass invested, in nutrient-rich environments (62–64). Levels of ho^5^U and mo^5^U also rise quickly during lag phase and then stabilize. Structurally, the hydroxyl group of ho^5^U stabilizes the anticodon loop in canonical Watson-Crick geometry, enabling pairing with U- and C-ending codons, whereas methylation to mo^5^U is required to stabilize the enol form of U to promote pairing with G-ending codons (65). Early increases in these modifications could therefore expand the decoding capacities of their respective tRNAs and support overall translational efficiency during rapid growth.

In many microorganisms, when nutrients begin to be depleted (on transition to stationary phase), carbon catabolite repression is lifted to enhance bacterial survival in the absence of preferred carbohydrates. During this time, Q modification levels decline and subsequent increases in unmodified Q-dependent tRNAs may then facilitate translation of less-biased and U-ending codon-biased transcripts, which are more abundant in proteins associated with metabolism of other (non-glucose) carbon substrates. m^2^A levels decrease in log phase and then increase through stationary phase. The fact that RlmN installs m^2^A in both tRNA and rRNA complicates understanding the role of the modification in the different RNA types. Plants, however, have separate m^2^A writers, permitting interrogation of the function of each (66). In *Arabidopsis*, m^2^A37 helps maintain a relaxed tRNA conformation and facilitates decoding of tandem m^2^A-bearing-tRNA-dependent codons (66). Although there is no reported preference that m^2^A bestows for one synonymous codon or another, Watson-Crick-pairing codons read by m^2^A-modifiable tRNAs are correlated with ribosomal and glycolytic proteins. Higher levels of m^2^A in stationary phase may stabilize codon-anticodon pairing in general, possibly promoting translational efficiency despite decreasing levels of ribosomes. Levels of s^4^U generally decrease over time. These trends are consistent with those previously reported in *Escherichia coli*, in which the fraction of s^4^U-modified tRNAs for select isoacceptors decreased with increasing growth rate (11). This modification plays a role in the intracellular stability of tRNAs, with hypomodification shown to result in rapid tRNA decay by the RNA degradasome (67). Decreased s^4^U levels in late stages of growth may reflect or contribute to translational slowdown.

It remains to be seen whether this observed growth phase-dependent tRNA reprogramming is unique to *E. faecalis* or is also present in other bacteria. *Vibrio cholerae* and *Escherichia coli* were also recently reported to have ribosomal proteins enriched in C-ending Q-dependent codons, despite a genome-wide preference for their U-ending counterparts (27). Regardless, the data presented here suggest a role for tRNA reprogramming and codon-biased translation in *E. faecalis* growth and proffer a framework on which future mechanistic studies, in this organism or others, can be designed. Furthermore, the comprehensive analyses and datasets provided herein will serve as a resource for the microbiology community.

## MATERIALS AND METHODS

### Bacterial growth conditions

*E. faecalis* OG1RF (68, 69) cultures were grown statically overnight (14-18 h) at 37 °C under aerobic conditions in BBL^TM^ Brain Heart Infusion broth (BHI, Becton Dickinson, 211059). For growth time-course experiments, overnight cultures of equivalent optical density (OD_600_ ∼ 1.5) were diluted 1:25 directly into fresh BHI media. Cultures were grown statically at 37 °C under aerobic conditions for 6 h, with absorbance measurements (OD_600_) recorded every hour.

### Bacterial lysis for RNA and protein extraction

Bacterial cultures (10-35 mL) were harvested by centrifugation at 5000*g* at 4 °C for 5 min. Cell pellets were then washed once with 35 mL of ice-cold PBS and pelleted again by centrifugation (5000*g* at 4 °C for 5 min). The wash supernatant was decanted, and the cell pellets were immediately resuspended in 1 mL of ice-cold TRIzol Reagent (Thermo Fisher, 15596026) and added to Lysing Matrix B tubes (MP Biomedicals, 116911100) on ice. Bacteria were lysed by bead-beating (3 x [40 s beat, 2 min rest on ice]) using a FastPrep-24 5G instrument (MP Biomedicals, 116005500) at a rate of 6.0 m/s. Samples were then centrifuged at 12000*g* at 4 °C for 5 min. The supernatants were transferred to new tubes to which 200 μL of chloroform was added. Samples were shaken vigorously for 15 s and then allowed to settle at ambient temperature for 2 min. The resulting mixtures were centrifuged at 12000*g* at 4 °C for 15 min, enabling separation of the aqueous (containing RNA) and organic (containing protein) phases.

### Large and small RNA isolation

Large and small RNAs were isolated from a portion (200 μL) of the extracted aqueous phase by solid-phase extraction using a PureLink miRNA Isolation Kit (Thermo Fisher, K157001) according to the manufacturer’s protocol with minor modification. Prior to transfer to the first cartridge, ethanol was added to the aqueous phase to a final concentration of 15% (v/v). This ethanol concentration was determined empirically to minimize retention of small RNAs on the first cartridge while still ensuring complete retention of large RNAs on the first cartridge. Ethanol was added to the flowthrough of the first cartridge to give a final concentration of 70% (v/v), prior to addition to the second cartridge. Samples on both cartridges were washed twice with wash buffer, spun dry, and then eluted in 50 μL pre-warmed (70 °C), nuclease-free (non-DEPC-treated) water. RNA concentrations were determined using a NanoDrop spectrophotometer, and RNA quality and purity was assessed by Agilent 2100 Bioanalyzer pico and small RNA microfluidic chips.

### RNA digestion and LC-MS/MS-based quantification of modifications

RNAs were hydrolyzed and dephosphorylated using a cocktail of enzymes, including benzonase, alkaline phosphatase, and phosphodiesterase I, at 37 °C for 6 h. Ribonucleoside analysis was performed using a Thermo Hypersil Gold aQ C18 column (100 × 2.1 mm, 1.9µm) on an Agilent 1290 Infinity II HPLC system with in-line UV detector coupled to an Agilent 6495 triple quadrupole mass spectrometer. Dynamic MRM was used for analyte detection and quantitation, with modification levels normalized to the sum of the UV signals (260 nm) for the unmodified nucleosides (A, U, G, C). Additional details are available in **Method S1**.

### Absolute quantification RNA-sequencing (AQRNA-seq)

To quantify tRNA isoacceptor levels, small RNA (75 ng) was used to prepare cDNA libraries for AQRNA-seq as previously described (56, 70), with targeted modifications to mitigate polymerase fall-off and primer dimer formation. Specifically, polymerase fall-off was minimized by replacing AlkB demethylation with a high-processivity reverse transcriptase (SuperScript IV; Thermo Fisher, 18091050), which enhances read-through across diverse RNA modifications and increases full-length cDNA yield. In addition, primer dimer formation was minimized by incorporating a post-reverse transcription digestion using *Escherichia coli* exonuclease I (ExoI, NEB, M0293S) to remove excess reverse transcription primers, thereby eliminating the need for gel purification. Libraries were quantified using a LightCycler 480 Real-Time PCR System (Roche), and library quality was assessed using the AATI Fragment Analyzer (Agilent). Pooled samples were cleaned using 0.9X volume PCRClean DX beads (Aline Biosciences) prior to 75 bp paired-end sequencing on an Illumina MiSeq platform (v3 reagent kit) with custom forward and reverse sequencing primers.(56, 70) A list of 37 mature, sequence-unique tRNAs present in *E. faecalis* OG1RF was curated from the Genomic tRNA Database (GtRNAdb). Raw sequences were processed using a custom AQRNA-seq analytical pipeline to yield tRNA abundances.

### Total RNA isolation

Total RNA was isolated from a portion (120 μL) of the extracted aqueous phase (see **Bacterial lysis for RNA and protein extraction**) using a RNeasy Mini Kit (Qiagen, 74106) according to the manufacturer’s protocol, including on-column DNase I treatment, with elution in 50 μL nuclease-free water. RNA quality was assessed using an Agilent 2100 Bioanalyzer pico microfluidic chip and RNA concentrations using a Qubit^TM^ RNA High Sensitivity Assay Kit (Thermo Fisher, Q32852) with a Qubit 2.0 fluorimeter. DNA contamination in the samples was determined to be < 10% using QuantiFluor® dsDNA and RNA Systems (Promega, E2671 and E3310, respectively).

### RNA-sequencing (RNA-seq)

Purified total RNA was processed using the Illumina Stranded Total RNA Library Preparation protocol (Standard, without rRNA depletion). Libraries were prepared using the IDT for Illumina RNA UD Indexes, and 150 bp paired-end sequencing was performed on an Illumina NextSeq 2000 platform using XLEAP-SBS chemistry. Raw sequenced reads were processed using standard open-source packages and tools for bacterial RNA-seq data processing. Briefly, sequence quality of the raw FASTQ files was assessed using FastQC (71), after which the paired-end reads were adapter and quality trimmed using fastp (72) with automatic paired-end adapter detection and a minimum post-trimming read length of 40 nt. Trimmed reads were aligned to the *E. faecalis* OG1RF reference genome (GenBank CP002621.1) using BWA-MEM (73) with default parameters. The resulting SAM output was converted to BAM using samtools (74), then coordinate sorted and indexed, and read counts per coding sequence (CDS) were generated from sorted BAMs using the featureCounts package (75). Per-sample counts attributed to CDS by locus tag values were finally assembled for downstream analysis.

### Protein isolation

Ethanol (300 μL) was added to the entire volume of the extracted organic phase (see **Bacterial lysis for RNA and protein extraction**), and samples were centrifuged at 2000*g* for 5 min at 4 °C to precipitate any remaining DNA. The supernatants were transferred to fresh tubes to which 2 volume equivalents of isopropanol were added. Samples were mixed well and allowed to stand at ambient temperature for 10 min. Samples were then centrifuged for 5 min at 12000*g* at 4 °C and the supernatant removed carefully by pipette. Protein pellets were washed three times with 2 mL of 95% (v/v) ethanol (in water) and once more with 100% ethanol, with centrifugation at 7500*g* for 5 min at 4 °C. Supernatants were removed and the protein pellets air dried at ambient temperature for 10 min.

### Quantitative proteomics

Protein pellets were digested with Lys-C and trypsin and peptides labeled using the TMTpro 18-plex Label Reagent set (Thermo Fisher, A52045). Combined samples were subjected to high-pH fractionation prior to mass spectrometric analysis. Chromatographic separation was performed using an EASY-nLC 1000 column (Thermo Fisher) coupled to an Orbitrap Eclipse Tribrid mass spectrometer (Thermo Fisher) using data-dependent mode. Raw mass spectra were searched using Sequest available in Proteome Discoverer (v3.1, Thermo Fisher) against *E. faecalis* OG1RF primary protein sequences retrieved from NCBI (NCBI Reference Sequence: NC_017316.1). Additional details are available in **Method S2**.

### Hydrogen peroxide measurements

Bacteria-produced hydrogen peroxide levels were determined using a method adapted from Boonanantanasarn et al. (76) and Gaca et al. (49), wherein hydrogen peroxide production is measured using a horseradish peroxidase-coupled fluorometric assay. Bacterial cultures from various growth phases were prepared as described above (see **Bacterial growth conditions**) and a volume of culture that would yield an OD_600_ of 0.5 if resuspended in 500 µL was harvested by centrifugation at 10000*g* for 2 min at ambient temperature. Cells were washed twice in assay buffer (50 mM sodium phosphate buffer, pH 7.4, freshly prepared) and resuspended in 500 µL assay buffer. 50 uL of cell suspensions were added to wells of an opaque 96-well microtiter plate. Equivalent volumes of a series of hydrogen peroxide standards (in assay buffer) together with an assay buffer blank control were also added to the plate. Reactions were initiated by addition of 50 uL of an Amplex UltraRed/HRP working solution to give final solution concentrations of 20 µM Amplex UltraRed Reagent (Thermo Fisher, A36006) and 0.2 U/mL horseradish peroxidase (Thermo Fisher, 31490) in assay buffer. Fluorescence was read in an Infinite M Plex plate reader (Tecan) with excitation at 490 nm and emission at 590 nm, with measurements taken after 15 min. Background fluorescence, determined using the blank control, was subtracted from all measurements and hydrogen peroxide levels calculated based on the standard curve. Cell aliquots from each growth phase were serially diluted and plated on BHI agar for colony enumeration to permit normalization of hydrogen peroxide levels to cell count (CFU).

### Data mining and statistical analysis

All data mining and statistical analyses were performed in either Python or R, unless otherwise indicated. Soft clustering of proteomics data was performed in R using the Mfuzz package (42). Raw protein intensities were normalized by total protein abundance per sample, normalized across samples by arithmetic mean, and then log2-transformed. Time-matched biological replicates were subsequently averaged, and protein z-scores determined prior to clustering. The optimal number of clusters was determined from a minimum centroid distance (*D_min_*) curve. Overrepresentation analysis was performed on the resulting protein clusters using Bioconductor’s clusterProfiler package and publicly available KEGG and in-house-curated GO databases (77–81). Raw RNA counts were normalized using DESeq2 (median-of-ratios normalization) (82), followed by variance-stabilizing transformation. Time-matched biological replicates were then averaged and transcript z-scores determined. Only proteins with corresponding transcripts (and vice versa) were included in calculations of the centroid values (**Fig. 2A**). R scripts and associated data used to generate **Fig. 2** are available at https://github.com/kline-unige/multiomics-clustering. Gene-specific and genome-based codon analytic measures were made using previously described methods (9). Each protein-coding gene was analyzed from start to stop codon and the number of each of 64 codons was counted. From these data, gene-specific isoacceptor codon frequencies and corresponding z-scores were computed. Raw tRNA abundances were normalized using DESeq2 (median-of-ratios normalization) in R (82).

## DATA AVAILABILITY

All source data have been deposited in public repositories: LC-MS/MS RNA modification data is available through the PRIDE repository accession XXXX (XXXXXX); AQRNA-seq data is available through GEO accession XXX (XXXX); RNA-seq data is available through GEO accession XXX (XXXX); and proteomics data is available through the PRIDE repository accession XXXX (XXXXXX). Processed data and all other supporting files are included in the

## Supplementary Information

## ACKNOWLEDGEMENTS

We thank Siok Ghee Ler, Jeremy Wang, and Radoslaw Sobota from A*STAR IMCB and the A-PROMPT research support center for guidance with proteomics analyses. We thank Justine Dacanay and the SCELSE sequencing facility for preparing and sequencing the RNA-seq libraries. We thank Stuart Levine and the MIT BioMicro Center for sequencing the AQRNA-seq libraries. We thank Cristina Colomer-Winter for advice on H_2_O_2_ assays. ChatGPT (OpenAI) was used to assist in coding for quantitative graphic generation in Python, with careful review, editing, and validation by authors. Pictorial graphics were made using Biorender.com and chemical structures drawn in MolDraw.

This work was supported by the National Research Foundation, Singapore, under its Campus for Research Excellence and Technological Enterprise (CREATE) program, through core funding of the Singapore-MIT Alliance for Research and Technology (SMART) Centre, Antimicrobial Resistance Interdisciplinary Research Group (AMR IRG). This work was partially supported by the Singapore Centre for Environmental Life Sciences Engineering (SCELSE), funded by the National Research Foundation and Ministry of Education, Singapore, under its Research Centre of Excellence Program. Additional support was provided to K.A.K. by the Singapore National Medical Research Council Open Fund (MOH-000645), the Société Académique de Genève, and the Swiss National Science Foundation (310030_219227); and to T.J.B. by the National Institutes of Health (R01GM143749 and R01ES026856).

## AUTHOR CONTRIBUTIONS

M.M.M., M.V., K.A.K., and P.C.D. conceived the study. M.M.M. performed growth experiments and isolated analytes with help from M.V. and K.T. M.M.M. performed proteomics experiments and analyzed data. C.M.A. processed RNA-seq data and implemented soft-clustering analysis of proteomics data, overrepresentation analysis, and comparison with transcriptomics. T.J.B. performed codon analytics. M.M.M., K.T., and I.H. acquired and analyzed RNA modification data. M.M.M. and K.T. performed hydrogen peroxide measurements. M.M.M. prepared AQRNA-seq libraries and analyzed data with technical input from R.C. M.R. and Y.Y. identified tRNA-modifying enzyme homologs and biosynthetic pathways in *E. faecalis*. A.D. synthesized a modified ribonucleoside standard. K.P. assisted in project administration. M.M.M., K.A.K., and P.C.D. wrote the manuscript with input from all authors.

