## Supplementary Information for "Growth-dependent tRNA Reprogramming and Codon Bias Link Translation to Metabolic State in *Enterococcus faecalis*"

**Supplementary Figures**

**Fig. S1.** Queuosine structure and biosynthesis.

**Fig. S2.** Hydrogen peroxide produced by *E. faecalis* during different growth phases.

**Fig. S3.** 5-hydroxyuridine and 5-methoxyuridine structures and biosynthesis.

**Fig. S4.** 4-thiouridine structure and biosynthesis.

**Fig. S5.** 2-methyladenosine structure and biosynthesis.

**Fig. S6.** Ribosomal RNA modification changes during growth.

**Supplementary Tables** (separate Excel files)

**Table S1.** Raw data and subsequent calculations/normalizations for *E. faecalis* OG1RF proteomics data across the growth time course.

**Table S2.** Mfuzz soft-clustering analysis of z-score-normalized proteomics data.

**Table S3.** Overrepresentation analysis of protein clusters using KEGG and GO databases.

**Table S4.** Raw data, subsequent calculations/normalizations, and cluster centroids for *E. faecalis* OG1RF transcriptomics data across the growth time course.

**Table S5.** Gene-specific codon usage, isoacceptor codon frequencies, and corresponding isoacceptor frequency-based z-scores for all *E. faecalis* OG1RF protein-coding genes.

**Table S6.** tRNA isoacceptor decoding of codons in *E. faecalis* OG1RF.

**Table S7.** Raw data, subsequent normalization, and $\log_2$(fold-change) calculations for *E. faecalis* OG1RF tRNA modification data across the growth time course.

**Table S8.** Raw data, subsequent normalization, and $\log_2$(fold-change) calculations for *E. faecalis* OG1RF tRNA isoacceptor data across the growth time course.

**Table S9.** LC-MS/MS detection parameters for RNA modifications identified in *E. faecalis* OG1RF.

**Supplementary Methods**

**Method S1.** RNA digestion and LC-MS/MS-based quantification of modifications.

**Method S2.** Quantitative proteomics.

**Supplementary Figures**

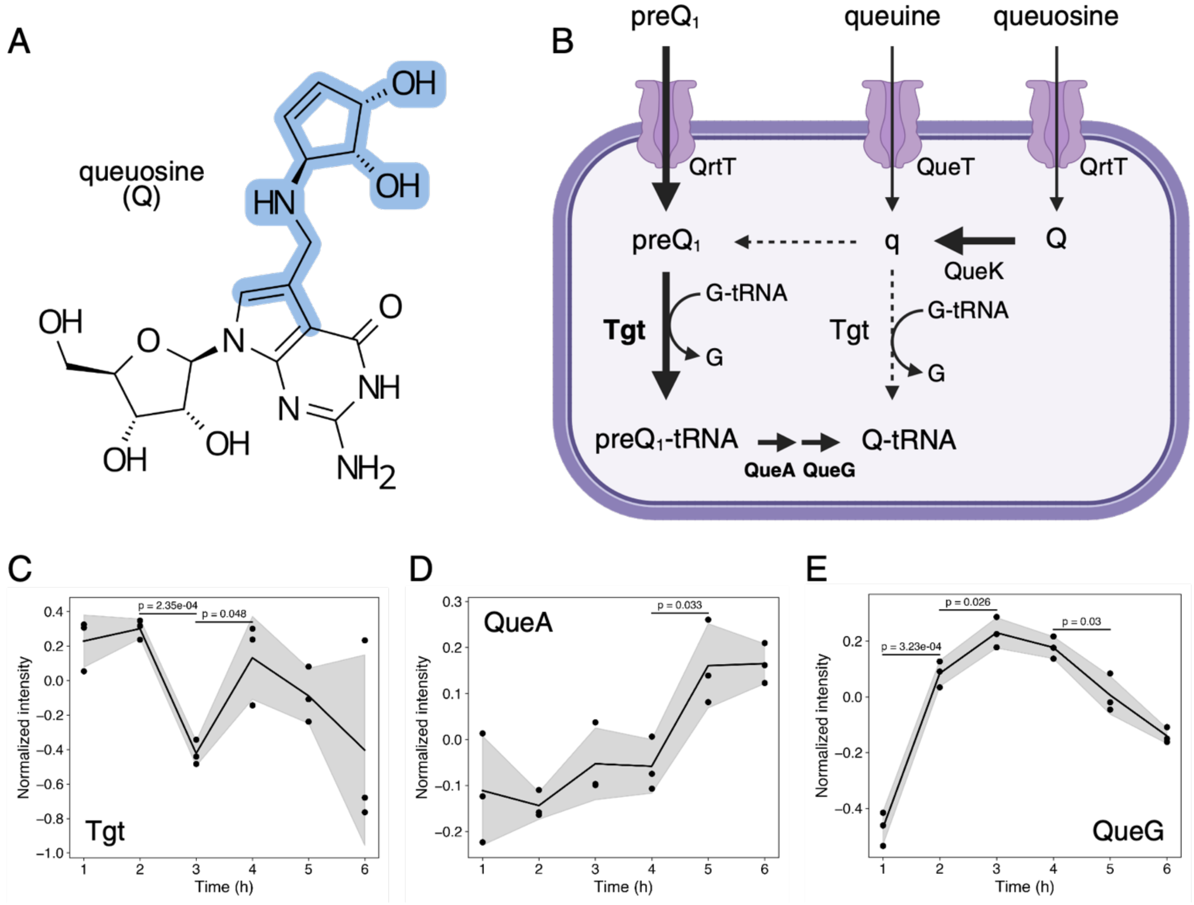


**Fig. S1. Queuosine structure and biosynthesis. (A)** Chemical structure of queuosine (Q), a modified derivative of guanosine, with the modification highlighted in blue. **(B)** Presumed pathway of Q biosynthesis in *E. faecalis* OG1RF based on homology. Bold arrows indicate the most likely pathway(s), narrow arrows less likely pathways, and dashed arrows instances of missing or absent enzymes in the organism. Levels of predicted biosynthetic enzymes (indicated in bold) are shown over time: **(C)** Tgt (OG1RF_RS03250), **(D)** QueA (OG1RF_RS03125), and **(E)** QueG (OG1RF_RS06700). Shaded regions represent the mean ± SD (n = 3 biological replicates). Protein levels at adjacent time points were compared using two-sided Welch's t-tests, with significant differences ($p < 0.05$) indicated by p-values on the graphs.

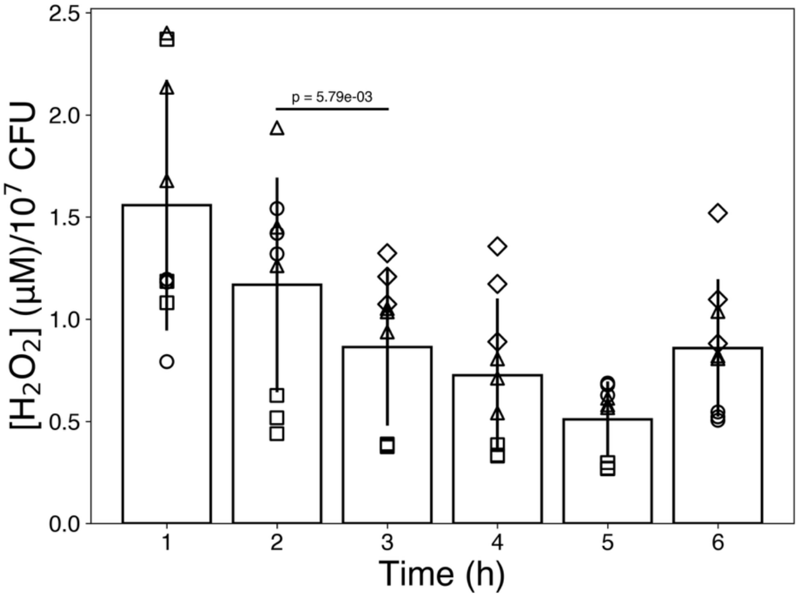


**Fig. S2. Hydrogen peroxide produced by *E. faecalis* during different growth phases.** Data represent four independent experiments (see different data markers/shapes) with three biological replicates for time points included in those experiments (each time point has three biological replicates from three independent experiments, i.e. nine data points total). Hydrogen peroxide ($H_2O_2$) levels were analyzed using linear mixed-effects modeling (time = fixed effect; experiment = random effect) and plotted as mean ± SD. Pairwise comparison of $H_2O_2$ levels at adjacent time points was performed and p-values adjusted using Benjamini-Hochberg FDR correction. Significant differences ($p < 0.05$) are indicated by p-value on the graph.

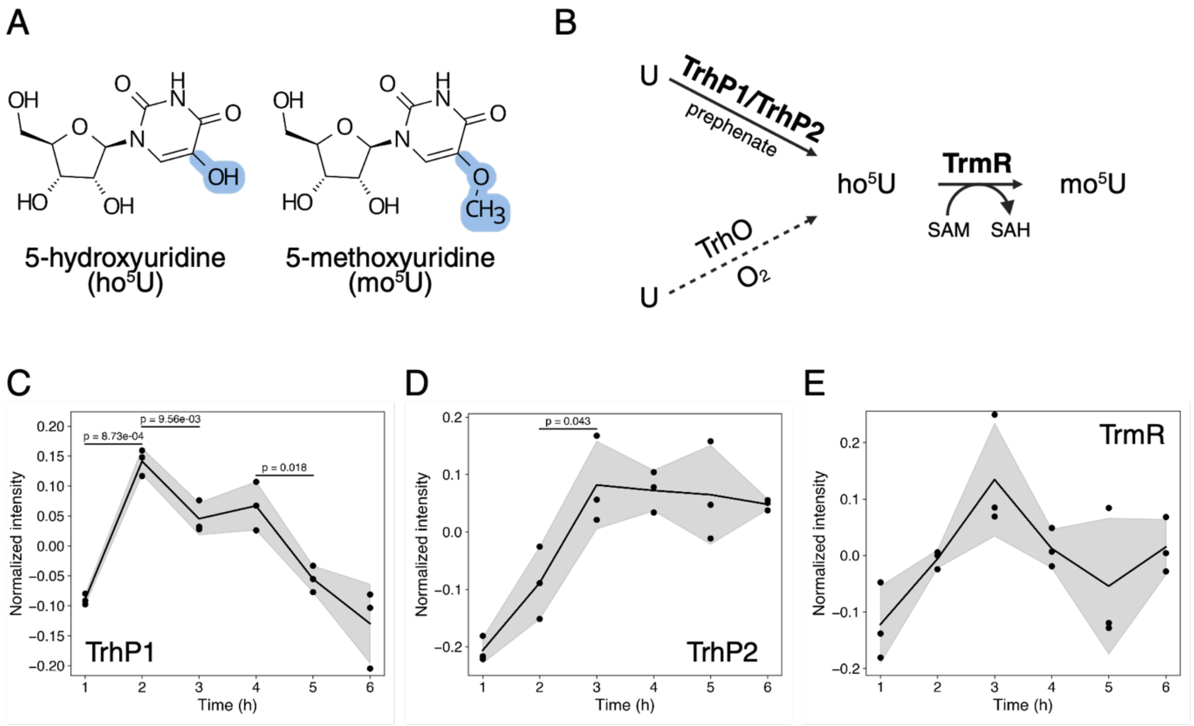


**Fig. S3. 5-hydroxyuridine and 5-methoxyuridine structures and biosynthesis. (A)** Chemical structures of 5-hydroxyuridine (ho$^5$U) and 5-methoxyuridine (mo$^5$U), uridine derivatives with modifications highlighted in blue. **(B)** Presumed pathway of ho$^5$U and mo$^5$U biosynthesis in *E. faecalis* OG1RF based on homology. Solid arrows indicate likely pathways, whereas the dashed arrow indicates a missing or absent biosynthetic pathway in the organism. Levels of predicted biosynthetic enzymes (indicated in bold) are shown over time: **(C)** TrhP1 (OG1RF_RS12885), **(D)** TrhP2 (OG1RF_RS12890), and **(E)** TrmR (OG1RF_RS10760). Shaded regions represent the mean ± SD (n = 3 biological replicates). Protein levels at adjacent time points were compared using two-sided Welch's t-tests, with significant differences ($p < 0.05$) indicated by p-values on the graphs.

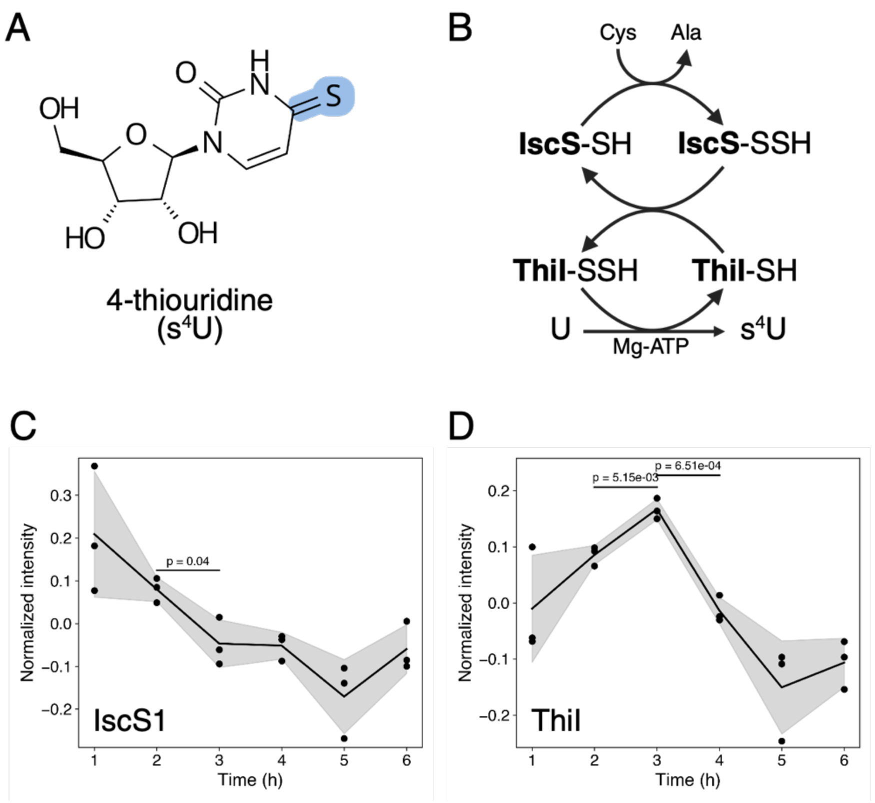


**Fig. S4. 4-thiouridine structure and biosynthesis. (A)** Chemical structure of 4-thiouridine ($s^4U$), a uridine derivative with modification highlighted in blue. **(B)** Presumed pathway of $s^4U$ biosynthesis in *E. faecalis* OG1RF based on homology. Levels of predicted biosynthetic enzymes (indicated in bold) are shown over time: **(C)** IscS1 (OG1RF_RS01435) and **(D)** ThiI (OG1RF_RS11450). Shaded regions represent the mean ± SD (n = 3 biological replicates). Protein levels at adjacent time points were compared using two-sided Welch's t-tests, with significant differences ($p < 0.05$) indicated by p-values on the graphs.

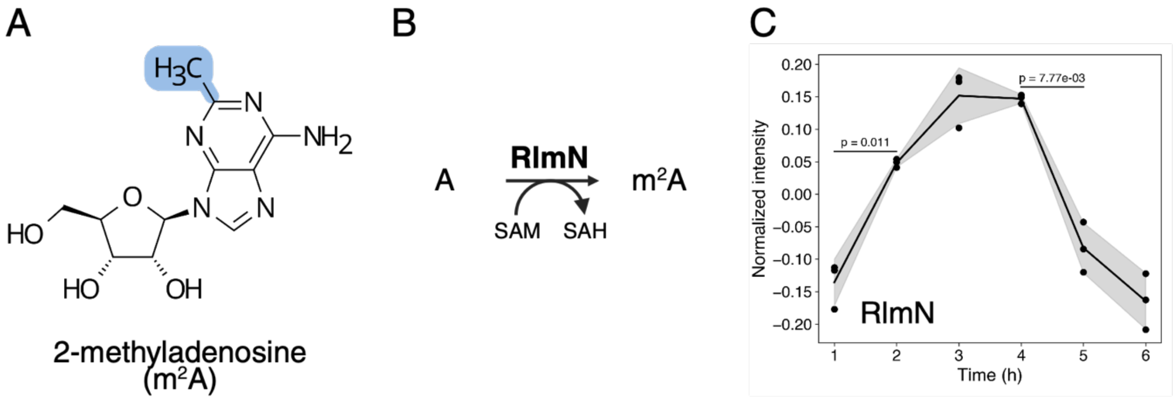


**Fig. S5. 2-methyladenosine structure and biosynthesis. (A)** Chemical structure of 2-methyladenosine, an adenosine derivative with modification highlighted in blue. **(B)** Established pathway of $m^2A$ biosynthesis in *E. faecalis* OG1RF. **(C)** Levels of the biosynthetic enzyme (indicated in bold), RlmN (OG1RF_RS08485), shown over time. Shaded regions represent the mean ± SD (n = 3 biological replicates). Protein levels at adjacent time points were compared using two-sided Welch's t-tests, with significant differences ($p < 0.05$) indicated by p-values on the graphs.

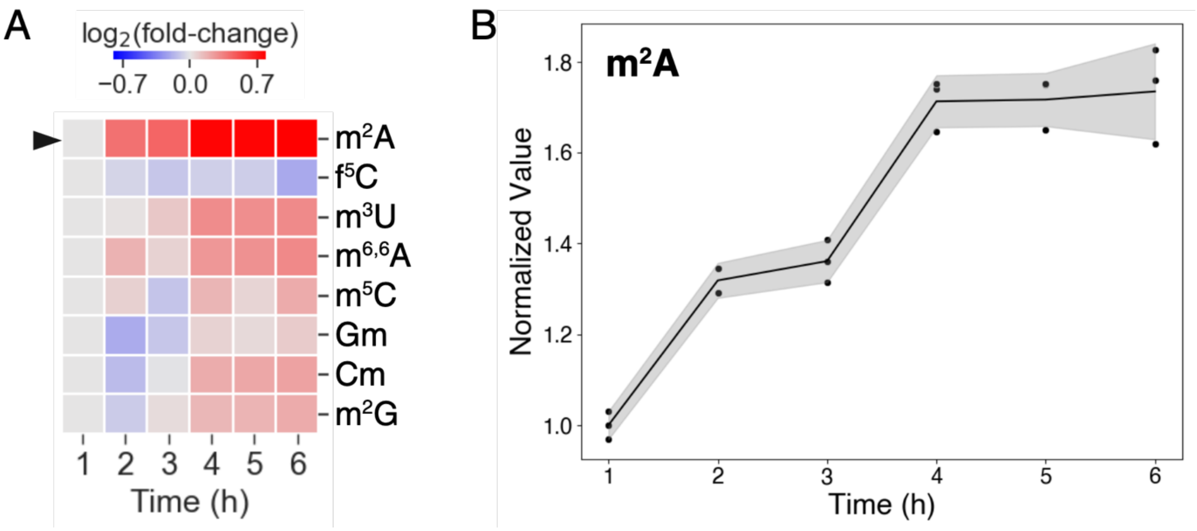


**Fig. S6. Ribosomal RNA modification changes during growth. (A)** Heatmap of large RNA (16S/23S rRNA) modification levels over the growth time-course, expressed as $\log_2$(fold-change) with respect to modification levels at t = 1 h. Values represent the average of three biological replicates. **(B)** Scatterplot of $m^2A$ levels (indicated by arrow in **(A)**) over time, expressed relative to modification level at t = 1 h. Shaded regions represent the mean ± SD.

**Method S1. RNA digestion and LC-MS/MS-based quantification of modifications.**

Large and small RNAs (500 ng – 1.25 µg) were digested with benzonase (0.25 U/µL, Sigma, E8263), alkaline phosphatase from bovine intestinal mucosa (0.1 U/µL, Sigma, P5521), and phosphodiesterase I (0.003 U/µL, Sigma, 3243) in a solution of 5 mM Tris (pH 8.0) and 2.5 mM $MgCl_2$ supplemented with 0.1 mM deferoxamine (antioxidant, Sigma, D9533), 0.1 mM butylated hydroxytoluene (antioxidant, Sigma, W218405), 0.1 µg/mL coformycin (adenosine deaminase inhibitor, NCI, 27781713), and 50 nM [$^{15}N_5$]-2'-deoxyadenosine (for monitoring triple quadrupole instrument performance; not for quantitation purposes, Cambridge Isotope Laboratories, NLM-3895) in a total volume of 25 µL. Samples were incubated at 37 °C for 6 h. The digestion mixtures were then centrifuged at 13000*g* at 4 °C for 10 min and the supernatants subjected to LC-MS/MS analysis.

Ribonucleoside analysis (25 – 200 ng on column) was performed using a Thermo Hypersil Gold aQ C18 column (100 × 2.1 mm, 1.9µm) on an Agilent 1290 Infinity II HPLC system with in-line UV detector coupled to an Agilent 6495 triple quadrupole mass spectrometer. Liquid chromatography was performed with a column temperature of 35 °C and a flow rate of 0.35 mL/min, with the following gradient: 0-4 min, 100% solvent A (0.1% (v/v) formic acid in water); 4–5.1 min, 0–1% solvent B (0.1% formic acid (v/v) in acetonitrile); 5.1-6.3 min, 1-6% solvent B; 6.3-7 min, 6% solvent B; 7-9 min, 6-50% solvent B; 9-11 min, 50-80% solvent B; and 11-15 min, 80% solvent B. The mass spectrometer was equipped with an Agilent Jet Stream electrospray ionization source and operated in positive mode with the following parameters: gas temperature, 120°C; gas flow, 11 L/min; nebulizer, 40 psi; sheath gas temperature, 400 °C; sheath gas flow, 12 L/min; capillary voltage, 1500 V; and nozzle voltage, 0 V. Dynamic multiple reaction monitoring (dMRM) was employed for analyte detection, with collision energies optimized for maximal sensitivity (**Table S9**). Modified ribonucleoside identities were confirmed by comparison of analyte retention times and MRM transitions to those of synthetic standards. Analyte peak areas were normalized to the sum of the UV signals (260 nm) for the unmodified nucleosides (A, U, G, C).

**Method S2. Quantitative proteomics.**

Protein pellets were resuspended in 8 M urea in 100 mM triethylammonium bicarbonate buffer (TEAB, pH 8.5) and protein amounts quantified by BCA assay. Protein samples (28 μg) were reduced with 10 mM tris(2-carboxyethyl)phosphine) for 20 min at 55 °C, followed by alkylation with 55 mM 2-chloroacetamide for 30 min at room temperature in the dark. Samples were then diluted with additional 100 mM TEAB (pH 8.5) to give a final urea concentration of 1 M. Protein was digested with 2.5 μg lysyl endopeptidase (Lys-C, FUJIFILM Wako, 129-02541) for 4 h at 37 °C with gentle agitation, followed by addition of 2.5 μg of sequencing grade modified trypsin (Promega, V5117) and incubation overnight at 37 °C with gentle agitation. Digestion efficiency was assessed and additional trypsin (1 μg) added with incubation overnight at 37 °C with gentle agitation. Digestion was quenched by addition of trifluoroacetic acid to a final concentration of 1% (v/v) and samples desalted using Oasis HLB cartridges (30 mg, Waters, WAT058951) with acidic buffers and the eluate dried to completion by vacuum centrifugal evaporator at 30 °C. Samples were resuspended in 35 μL of 100 mM TEAB (pH 8.5) and peptides quantified by Quantitative Fluorometric Peptide Assay (Pierce, 23290). Tandem Mass Tag (TMT) labeling was performed on 10 μg peptides from each sample by addition of 4 μL of 25 μg/μL TMT label (in anhydrous acetonitrile) from the TMTpro 18-plex Label Reagent set

(Thermo Fisher, A52045). Labeling efficiency was assessed and labeling reactions quenched by addition of 60 µL of 1 M ammonium formate (pH 10) for 5 min at room temperature. Labeled peptide samples were combined and dried to completion by vacuum centrifugal evaporator at 45 °C, followed by resuspension in 0.1% (v/v) formic acid in water. Acidic desalting was performed using Oasis HLB cartridges (60 mg, Waters, 186000679) and the eluate dried to completion by vacuum centrifugal evaporator at 45 °C. Samples were resuspended in 10 mM ammonium formate (pH 10) and separated into seven fractions by high-pH fractionation using spin columns prepared with Dr. Maisch ReproSil-Pur Basic-C18 resin (80 mg, IT Technologies Pte Ltd, r10.b9.0025). Fractions were dried via vacuum centrifugal evaporator and washed three times with 350 µL of 60% acetonitrile in water with 0.1% (v/v) formic acid to remove volatile salts (dried to completion each time). Dried fractions were resuspended in 10 µL of 0.1% (v/v) formic acid in water and 2 µL of each was injected for mass spectrometric analysis.

Chromatographic separation was performed using an EASY-nLC 1000 column (Thermo Fisher) coupled to an Orbitrap Eclipse Tribrid mass spectrometer (Thermo Fisher) using data-dependent mode. Peptides were separated using a 0-40% (v/v) acetonitrile gradient over 75 min, increasing to 83% over the next 8 min, and finally to 100% over 7 min, running at a constant flow rate of 300 nL/min. The following parameters were set for MS data acquisition: automatic gain control (AGC) set at standard with orbitrap resolution at 60,000 ranging from m/z 350 to 1550. Dynamic exclusion for precursors selected for fragmentation was set to 60 s. For fragmentation, the MS2 isolation window was set to 1.2 *m/z* of the selected precursor masses and higher-energy collision dissociation (HCD) was activated at 42% normalized collision energy. Fragment signals (MS2) were analyzed by orbitrap analyzer set to a resolution of 30,000 with the TurboTMT option selected for TMTpro reagents. The AGC target was customed at 150% and maximum injection time (IT) at 54 ms with centroid data type. Raw mass spectra were searched using Sequest available in Proteome Discoverer™ (v3.1, Thermo Fisher) against *E. faecalis* OG1RF primary protein sequences retrieved from NCBI (NCBI Reference Sequence: NC_017316.1). The mass tolerance for precursors and fragments was set to 10 ppm and 0.06 Da, respectively; and TMT quantitation was based on reporter ion quantifier node set at 20 ppm for HCD. The target FDR was set at 0.01. Proteins identified with high-confidence (FDR = 1%) in all samples were used in downstream analyses.
